# Widespread transcriptional responses to the thermal stresses are prewired in human 3D genome

**DOI:** 10.1101/728220

**Authors:** Xiaoli Li, Bingxiang Xu, Xiao Li, Danyang Wang, Ziyang An, Yan Jia, Jing Niu, Juntao Gao, Michael Q Zhang, Feifei Li, Zhihua Zhang

## Abstract

Temperature changes is one of the most common environmental stress that consequences with massive phenotypic responses for almost all the life forms. The dysregulation of heat shock (HS) response genes had been found associated with various severe diseases, including cancer. Although the HS response has been well studied in animal cells, it remains elusive whether or not the cells response to cold shock (CS) similarly. Here, we comprehensively compared the changes of gene expression, epigenetic marks (H3K4me3 and H3K27ac), binding of genome architecture proteins (CTCF, SMC3 and Pol II) and chromatin conformation after HS and CS in human cells. Widespread expression change was observed after both HS and CS. Remarkably, we identified distinguished characters in those thermal stress responded genes at nearly all levels of chromatin architecture, *i.e*, the compartment, topological associated domain, chromatin loops and transcription elongation regulators, in the normal condition. However, the global chromatin architecture remains largely stable after both CS and HS. Interestingly, the thermal stresses responded genes are prone to spatial clustering even before the temperature changes. Our data suggested that the transcriptional response to the thermal stresses maybe independent to the changes of the high-level chromatin architecture, e.g., compartments and TAD, while it may be more dependent on the precondition of the chromatin and epigenetic settings at the normal condition.

## Introduction

Life always suffers various stresses from environment fluctuation, and how life responses to those fluctuation is critical to survival. Temperature change is one of the most common environmental stresses, it can be either higher (heat) or lower (cold) than one’s comfort temperature. The thermal stress can have broad effect to a range of important phenotypes, such as gestation and sex determination. There are tremendous diverse mechanisms to deal with the temperature changes in nature, from the macroscopic behavers, e.g. hibernation, to the molecular level reactions, e.g., gene express change (Carey et al., 2003; Sonna et al., 2002). There is a whole category of stress response genes which have been evolved to deal with various endogenous and exogenous stress (Fujimoto and Nakai, 2010; Kregel, 2002). For thermal stress, however, much attention has been paid to the study of the heat shock (HS) response in animals (Velichko et al., 2013). For example, whole genome transcriptional re-modulation has been observed in yeast (Chen et al., 2003), worm (Ni et al., 2016), fly (Duarte et al., 2016) and human cells (Lyu et al., 2018; Mahat et al., 2016b; Vihervaara et al., 2017) in response to HS. There are small group of genes which have been up-regulated in HS, and most of them were found associated with the heat shock protein (HSP) family, which probably is one of the most well studied protein family (Fujimoto and Nakai, 2010; Kregel, 2002), and it has been found play important roles in carcinogenesis (Wu et al., 2017). Cold shock (CS) response is also important to animals, e.g., hypothermia can be lethal (Sonna et al., 2002). And it has become growing important to understand the CS response of human cells as more organ transplants take place in modern medicine practice, while proper preserve organs in cold condition is an essential step to successful transplant (Al-Fageeh and Smales, 2006). However, the molecular mechanisms of CS response have been much understudied. Microarray data had been shown that there may also genome wide transcriptional alteration in response to CS in mammals (Beer et al., 2003), and a few cold shock proteins, such as CIRP and Rbm3, and HSP family may be involved in regulation of CS response (Carey et al., 2003; Fujita, 1999).

Gene transcription regulation has been associated with the chromatin spatial structures (Hubner et al., 2013). The alteration of chromatin spatial structure, e.g, the genome compartments (Lieberman-Aiden et al., 2009b), the topological associated domains (TAD) (Dixon et al., 2012), and chromatin loops (Rao et al., 2014), has been suggested underling the transcriptional responses to HS in multiple species. In yeast, up-regulated HSP genes undergo dynamic alteration in their 3D genome structure (Chowdhary et al., 2017; Chowdhary et al., 2019). In Drosophila, HS induces relocalization of architectural proteins from TAD borders to inside TADs, and causes a dramatic rearrangement in the 3D organization of chromatin (Li et al., 2015). In mammalians, the picture is much less clear. It has been shown that chromatin loops experience dramatic changes in human ES cells in response to HS, and the frequency of loop interactions is correlated with the level of nascent transcription (Lyu et al., 2018), while a recent preprint manuscript claimed largely stable chromosome architecture after short term HS in human myelogenous leukemia cell line (K562) (Ray et al., 2019). Alternatively, the regulation on the pausing Pol II has also been proposed (Vihervaara et al., 2018). However, the discussion on chromatin changes in CS is almost completely lack, which pose the comparison between CS and HS an open question.

Using ChIP-seq and in situ Hi-C technologies, we profiled multiple epigenetic marks (H3K4me3, H3K27ac and Pol II), binding of genome architecture proteins (CTCF and SMC3) and chromatin conformation in human K562 cells under normal (NM, 37°C), HS (42°C) and CS (4°C) conditions for 30 minutes. In addition to the genome-wide repression in gene transcription, there are remarkable number of genes be up-regulated in response to the thermal stresses, and many of them are not necessarily associate with HSP family. Surprisingly, comparing to NM, the chromatin architecture of human cells remains largely stable after both CS and HS, and even topical dynamic of chromatin has little association with this tremendous changes of transcriptome. Considering human being belongs homothermal animal, we reasoned that there should be more plans encoded in the genome for the thermal stress. Indeed, at NM condition, comparing to the down-regulated genes in response to thermal stress, the up-regulated genes have more binding of chromatin architecture proteins and transcription elongation regulators, the genome is more accessible and involves more chromatin loops to distal regulatory elements. Moreover, the thermal stress responded genes were found prone to spatial clustering in NM. Based on these data, we proposed that the genome response to the thermal stress is prewired in the architecture of chromatin as well as the landscape of epigenetic marks. The results provide a comprehensive comparison of how HS and CS will affect the genome, its architecture and its activity, and offered a reference resource for the study of environmental stress.

## Results

### Genome-wide transcriptional response to the thermal stresses are similar

To capture the live transcriptional response to the thermal stresses, we performed ChIP-Seq of Pol II in K562 cells under the normal (NM, 37°C), heat shock (HS, 42°C 30min) and cold shock (CS, 4°C 30min) conditions. The transcription level were quantified by the ChIP-seq reads per million of Pol II (CPM) (Amat et al.). The accuracy of CPM in quantifying live transcription were first assessed by super long genes analysis (i.e., > 150kb, (Mahat et al., 2016b)). If CPM is a fine index representing live transcription, we shall expect it remains intact after thermal stress in the region between 100kb to the transcription termination site (TTS) of the super long genes. This is because the transcript speed is about 3.1 Kb/minute in human (Wada et al., 2009), the transcript waves travels about 90Kb in 30 minutes, the Pol II signal after 100kb in the super long genes should be irrelevant to any interruptions by the thermal stresses. Indeed, using CPM we identified more (p=1.23e-40 and 2.42e-32 in CS and HS, respectively, χ^2^ test) differentially expressed normal genes (35.16% and 25.88% in CS and HS, respectively) than differentially expressed 50kb 3’end regions of super long genes (21.73% and 11.83% in CS and HS, respectively). This pattern can also be seen with in the super long genes. For example the CPM are dramatically altered in the 5’ end while remains largely intact in the 3’ends in response to the thermal stresses (Figure 1A, Figure S1A). Second, CPM is highly correlated with the precision nuclear Run-on Sequencing (PRO-seq) data in K562 cells. PRO-seq (Mahat et al., 2016a)is a technology that directly measures nascent transcription level (Mahat et al., 2016a). Comparing with the published PRO-seq data (Mahat et al., 2016c), the Spearman’s rank correlation coefficients (SpCC) between CPM and PRO-seq are, SpCC = 0.86, p < 2.26e-11 and SpCC= 0.84, p < 2.26e-11 in NM and HS, respectively. Moreover, the direction of gene expression changes determined by CPM and PRO-Seq (the sign of log2 fold changes) are well consistent with each other in HS (χ^2^ = 2153.04, p < 2.2e − 26, Figure S1B). Last, by applying the library size control scheme described in (Mahat et al., 2016b; Vihervaara et al., 2017), we identified 2492 and 3884 genes were up and down regulated in HS, respectively (fold change > 1.25 and p<0.001, Figure 1B). Among them, 537 out of 2492 (21.54%) and 2240 out of 3884 (57.67%) were also reported up and down regulated genes in PRO-seq data (Mahat et al., 2016c), respectively. Although, because the limited sensitivity and power of the CPM with Pol II ChIP-seq data, only 69.02% (537/778) and 36.60% (2240/6121) PRO-seq identified up and down genes were reported by the CPM analysis, the regulatory direction for the genes are consistent between two methods (Figure S1C). Moreover, the PRO-seq and ChIP-seq identified regulated genes are significantly overlapped between each other (p < 1e-4 for both up and down regulated genes, permutation test, Figure S1D). These results suggest that the CPM of Pol II ChIP-Seq can be a fine agent for gene transcript analysis in thermal stress.

**Figure 1.**
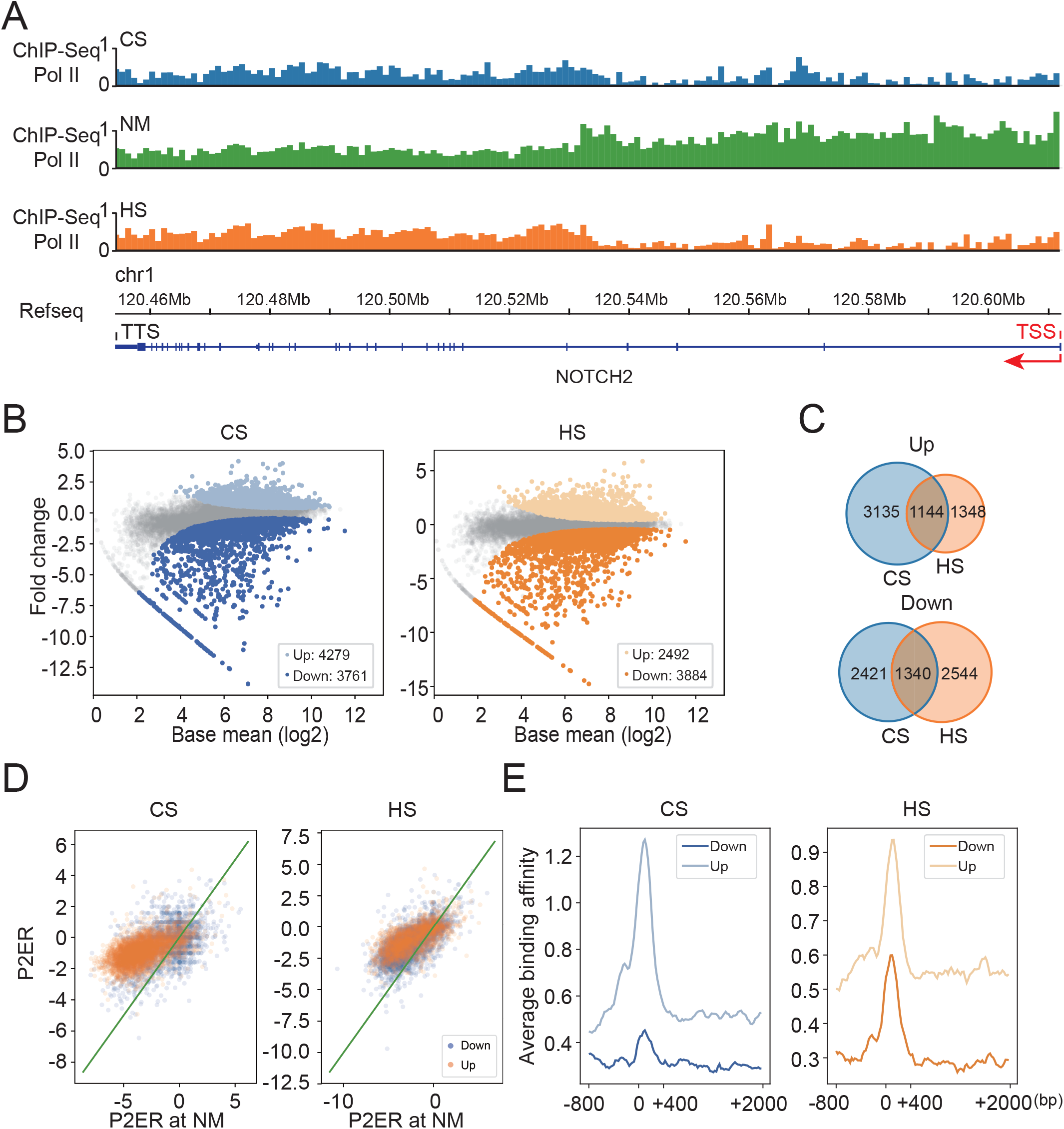
Genome-wide transcriptional response to the thermal stresses are similar. (A) After thermal stress, the Pol II are dramatically altered in the 5’end while remains largely intact in the 3’ends of super long genes. The example showed the distribution of Pol II ChIP-seq data in a downregulated gene NOTCH2. (B) MA plots of CPMs in CS and HS. Each dot represents a gene, and the light, grey and dark color represent up, null and down regulation, respectively. (C) The venn diagram shows the overlapping of up and down regulated gene in HS and CS. (D) The comparison of pause to elongation ratio (P2ER) between NM and the thermal stresses conditions. (E) The comparison of average NELF binding affinity in NM condition between the top up and down regulated genes (1115 ones (about 5% of all genes) with largest log2 fold changes), in response to the thermal stresses. 0 represents the TSSs.

Genome wide transcriptional response was observed in both CS and HS conditions, and the up and down regulated gene set is similar between the two stresses. First, genome wide transcriptional response was observed in both CS and HS conditions. There were 15.94% (3761 out of 23594) and 16.46% (3884) of RefSeq genes be identified as down-regulated, in CS and HS, respectively. There were 4279 and 2492 genes were found up-regulated in CS and HS, respectively (Figure 1B). Strikingly, a large portion of thermal responded genes are identical between HS and CS, i.e., 1340 and 1144 common down and up regulated genes, respectively (enrichment p-values < 1e-4, permutation tests for both condition, Figure 1C). For example, there are 29.41% and 47.05% genes in the heat shock protein (HSP) gene family been up regulated after CS and HS, respectively, and 18.42% of all HSP genes are shared up regulated between HS and CS (p=0.0001, permutation test). However, many of up-regulated genes are not associated with HSP family. In addition, Gene ontology (GO) analysis showed that the thermal stress induced genes, i.e. up regulated genes, enriched significantly similar GO terms between HS and CS (Figure S2A, Benjamini-Hochberg adjusted p < 0.05). There are 9 categories shared between HS and CS in their top 36 enriched categories (Figure S2). These two observations imply the cells may response to thermal stresses with similar transcription alteration.

It has been suggested that the mechanism of widespread transcription repression in HS is the blocked pause-release (Mahat et al., 2016c), as evidenced by the observation that, in the NM condition, the ChIP-seq peaks of Negative Elongation Factor (NELF) is enriched in the promoter region of HS up-regulated genes (Vihervaara et al., 2017; Yamaguchi et al., 1999). NELF is the factor that co-localizes with the paused Pol II to inhibit its release into productive elongation. Presumably, the enriched NELF blocked Pol II release from pausing, and the unbinding of NELF releases Pol II after HS. We wonder whether this is also the case in CS. To examine this speculation, we quantify the strength of Pol II releasing as the following (Vihervaara et al., 2017). The concentration of pausing and productive elongating Pol II were roughly inferred as the maximal ChIP-seq signal of Pol II in the region of [-100bp, 400bp], and [500bp, 1000bp], as oriented at TSS, respectively. This pause to elongation ratio (P2ER) is therefore represents the rate of pause releasing (Vihervaara et al., 2017).

Surprisingly, the P2ER were genome wide increases in response to the thermal stresses, and there were even 65.94% and 80.32% genes that being identified as down-regulated were also found has increased P2ER, in CS and HS, respectively (Figure 1D). This remains true even if we took a stricter definition of P2ER (Figure S1E). A plausible explanation to the observation of a large portion of down-regulated genes also have increase P2ER is that the increasement of P2ER in down-regulated genes may be smaller than that in the up-regulated genes (Figure S1E, p=1.26e-5 and 1.84e-279 in HS and CS, respectively, t test). Moreover, using the same ENCODE ChIP-seq data set as a previous study did (Consortium, 2012; Vihervaara et al., 2017), we also observed the enrichment of NELF-E at the promoters of the CS upregulated genes (Figure 1E). In fact, for the significantly regulated genes, the binding affinity of NELF-E in NM (TSS+100 to TSS+400) is positively correlated with the strength of gene regulation, i.e., log2 fold changes of CPM, in thermal stress conditions, i.e., SpCC = 0.11 and 0.20, p =2.01e-18 and 4.92e-76 for HS and CS, respectively (Figure 1E). Taken together, the alteration of pause-release may also connect to the transcriptional response to cold shock. However, more investigation-is needed to further confirm this speculation.

### Global chromatin 3D architecture remains largely stable in response to the thermal stresses

We asked if there are global chromatin 3D architecture changes in response to the thermal stresses in K562 cells. First, we examined the nucleus volumes of K562 cells in the thermal stressed conditions by confocal imaging, and found little significant changes of nuclear radius in response to the stress (p=0.2264 and 0.2628, for CS and HS, respectively, Mann Whitney U test, Figure 2A and B). Then, we performed in-situ Hi-C in K562 cells under the three conditions with 4, 2 and 3 replicates in CS, NM and HS, respectively. About 400 to 500 million valid contact read pairs are obtained in each conditions (Table S1), with high stratum adjusted correlation coefficient (SCC) between biological replicates (Yang et al., 2017), i.e. average SCC = 0.9874, 0.9984 and 0.9840 in NM, HS and CS, respectively. To further assess our Hi-C data quality, we calculated the Pearson’s correlation coefficient (PCC) in every 5Mb sliding windows comparing our in-situ Hi-C data in NM with the public data in Rao et al (Rao et al., 2014). The reason why we did comparison in the sliding windows was the K562 cells in different labs may carrying lab specific genome structure variations (Figure S3), since K562 has independently evolved about 44 years in the labs (Lozzio and Lozzio, 1975). We found the median PCC is 0.967 and more than 84% of the windows have PCCs higher than average 0.95 (Figure S4A). Indicating that the Hi-C data we generated is high quality. The Hi-C profiles are almost identical between the three conditions. At 50kb resolution, the Hi-C contact matrices of CS and HS are both highly correlated with NM (SCCs between NM and CS, and NM and HS are 0.95 and 0.98, respectively. Figure 2C and Figure S4B). Further, the contact frequency decay curves (P(s)) in the two thermal stress conditions are also identical to NM, i.e., the Jensen–Shannon divergences (JSD) between NM and thermal stress conditions are 9.27E−5 and 1.5E−4 for CS and HS, respectively (Figure 2D). These JSD is comparable to the JSD between biological replicates (mean JSD = 1.48E−4) in NM. Furthermore, little alterations of relative positions of chromosomes could be detected in response to the thermal stress. The relative positions of chromosomes were assessed by inter-chromosomal Hi-C contact reads frequency at 1Mb resolution. The inter-chromosomal contact frequencies were almost linearly correlated between NM and thermal stress conditions, i.e., mean PCC are 0.64 (from 0.46 to 0.87) and 0.66 (from 0.45 to 0.87) in HS and CS, respectively. The PCC level is comparable to the values between two biological replicates (mean 0.63, varying between 0.33 to 0.93, Figure 2E). Together, the global chromatin 3D organization, including the nuclear size, p(s) and relative positions of chromosomes, remains intact in response to thermal stress.

**Figure 2.**
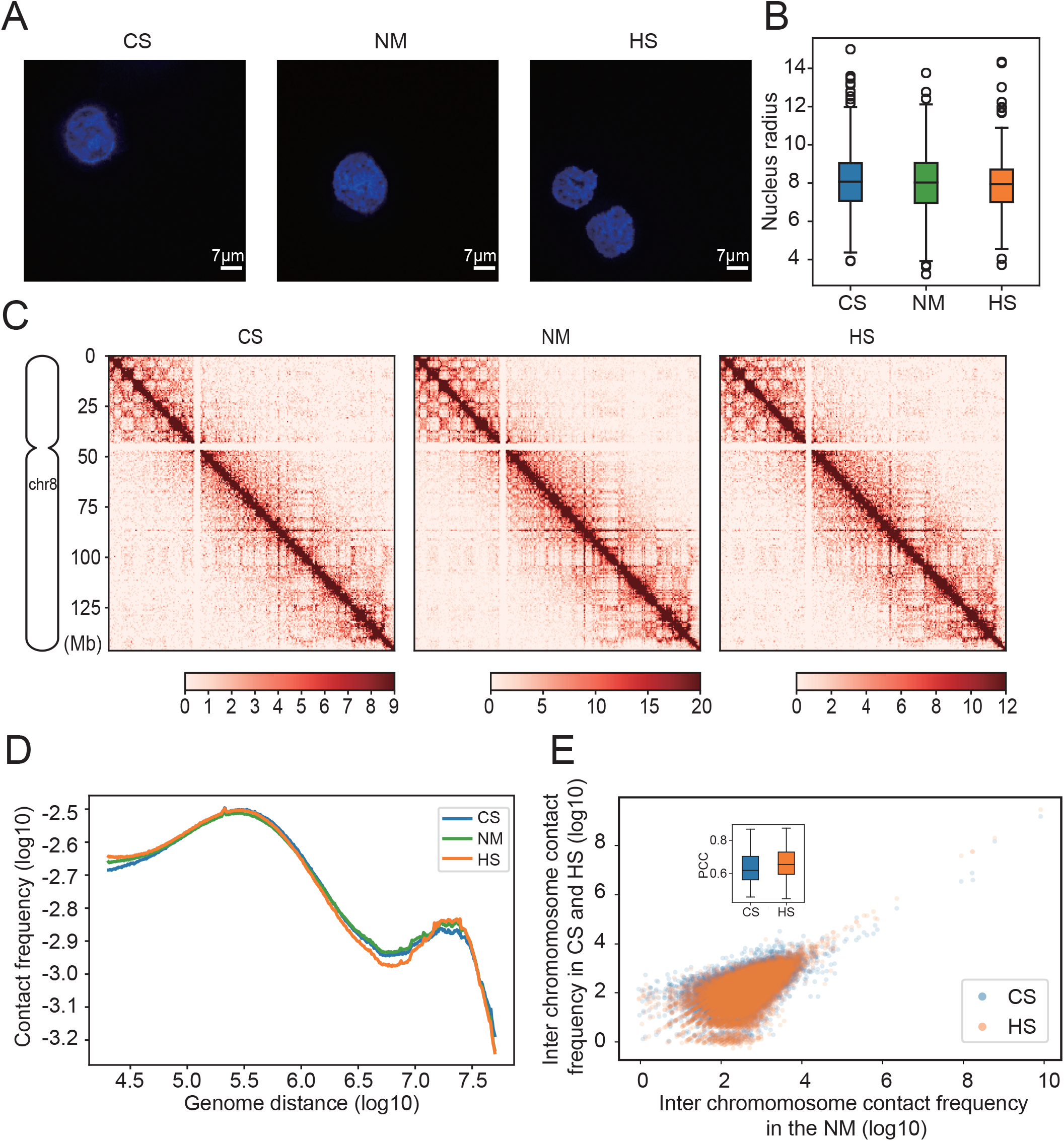
Global nucleus 3D organization remains stable in responses to the thermal stresses. The image example in (A) shows that the nucleus size remains stable after the thermal stresses in K562 cells, the distribution of 921 nucleus radius, 339, 299, 283 in CS, HS and NM, respectively, are showed in (B). The Hi-C contact matrices are highly correlated between the conditions. A representative example on chr8 at 100Kb resolution are showed in (C). (D) The contact frequency decay curves of Hi-C data at the thermal conditions. (E) The inter chromosome interaction are remains stable after the thermal stresses. The scatter plot represents an example of correlation of inter-chromosome contact between NM and the thermal stressed conditions. Each dot represents the Hi-C reads frequency between chr6 and chr8 in a 1Mbp length bin. The embedded boxplot shows the distribution of PCC between all chromosomes.

### Chromosomal Compartment changes are independent to transcriptional response to the thermal stresses

We next asked if the observed transcriptional response to the thermal stresses is reflecting the dynamic changes in chromosomal compartments, as the compartments have been associated with gene expression in many cell types (Lieberman-Aiden et al., 2009a). Surprisingly, the transcriptional changes were found to be independent from compartment changes in our data. First, the overall pattern of compartments is almost identical among the NM, HS and CS at 100Kb resolution (Figure 3A), although the scale of auto correlation coefficients shrunk after temperature stress. The first eigenvectors are almost linearly correlated between the three conditions (Supplemental Figure S5B). For example, in chromosome 8, the correlation coefficients are 0.97 and 0.98 between NM and CS and HS, respectively. 95.09% and 96.40% of the genome regions did not change compartment after CS and HS, respectively (Figure S5A). As a well-accepted mark for transcriptional actively that enriched in compartment A (Creyghton et al., 2010; Lieberman-Aiden et al., 2009a), H3K27ac were found largely unchanged in response to the thermal stress (The SpCC between NM and CS and HS are 0.88 and 0.89, respectively), which further evidenced stable compartments (Figure 3A). Second, a large portion of the genome regions that indeed switched their compartments are common between CS and HS (Figure S5A). At 100Kb resolution, there are 174 bins (18%=174/980 and 57%=174/303) experienced A to B compartment switch in both CS and HS, respectively, and 159 bins (38% = 159/422 and 22% = 159/724) experienced B to A compartment switch in both CS and HS, respectively. Second, the transcriptional responses to the thermal stresses is not directly associated with compartment switches, as the compartment switched region were not enriched for either up- or down-regulated genes, i.e., only 26 out of 2492 (p > 0.5, permutation test) and 43 out of 4279 (permutation test p > 0.5) up-regulated genes were overlap with those compartment B to A switched regions in HS and CS, respectively. For the down-regulated genes, only 1.62% and 2.55% experienced A to B compartment switch, in HS and CS, respectively. Together, the compartment switching and transcriptional response to the thermal stresses may be independent.

**Figure 3.**
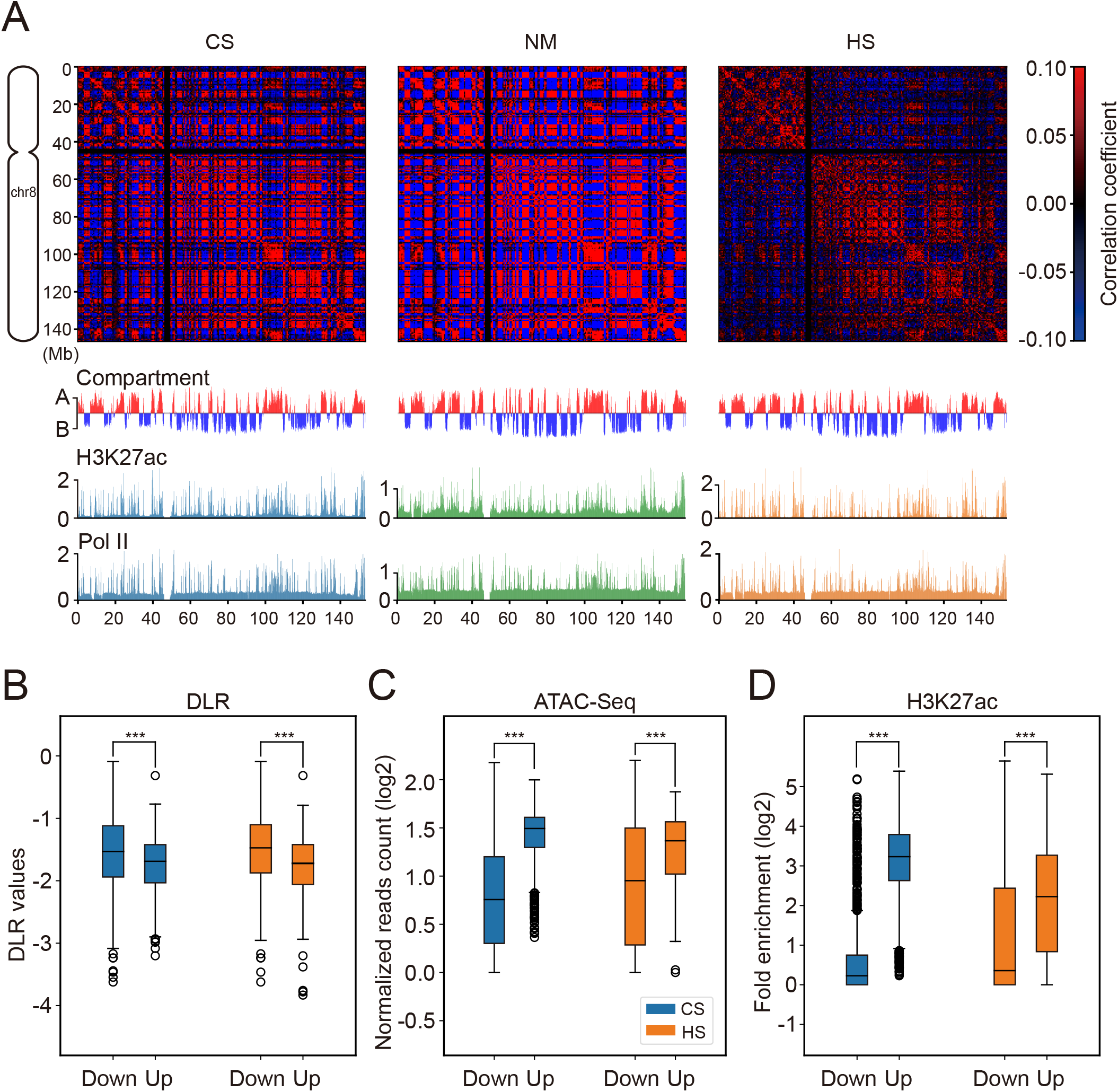
The responses of chromosomal compartment and gene expression to the thermal stresses are independent. (A) The auto-correlation matrix of Hi-C data at the three thermal conditions of chr8 at 100Kb resolution. The compartments are showed in the plaid patterns as well as the profile of the eigenvector 1. The comparison of DLR, ATAC-seq and H3K27ac ChIP-seq data between up and down regulated promoters, are showed in panel (B), (C) and (D). *** represents P < 0.001, Mann Whitney U test.

To further investigate the association between chromosome compartments and transcription responses to the thermal stresses, we examined the landscapes of chromatin accessibility and epigenetic marks (Fortin and Hansen, 2015). Strikingly, we found that the landscape of chromatin accessibility in NM can largely define how the loci response to the thermal stresses in transcription. The accessibility were inferred by the distal local ratio (DLR) of Hi-C data (Method, and (Heinz et al., 2018)). DLR is a fine index for accessibility, as it successfully distinguished the compartments at all three conditions, i.e. lower DLR values indicating compartment A (Figure S5C). In NM condition, the DLR of the 1115 top up regulated TSS are significantly smaller than that of 1115 most down-regulated TSS (p = 2.17e-18 and 6.83e-30, for CS and HS, respectively, Mann Whitney U test, Figure 3B). This is also true when the accessibility was inferred by ATAC-seq data, i.e., using only fragments shorter than 101bp (Figure 3C). The enrichment of ATAC-seq peaks in the promoter region of the top up regulated genes are significantly higher than those of the top down regulated genes after both CS and HS (P = 8.23e-165 and 1.30e-37 in CS and HS, respectively, Figure 3D and S5E). Meanwhile, the DLR of up and down regulated genes are indistinguishable between NM and HS (after removal of 11 outliers those who have DLR values close to zero at NM. p= 0.038, Mann Whitney U test), and only a slightly observable difference between NM and CS (after removal of 11 outliers, p=0.0042, Mann Whitney U test). The similar patterns were also found in H3K27ac ChIP-Seq data at NM (Figure 3D). The strength of H3K27ac in the promoter regions (defined as TSS-2Kb to TSS+2Kb) of the top up regulated genes are significantly higher than those of the top down regulated genes in NM condition, for both CS and HS (Mann Whitney U test, p = 1.09e-278 and 2.40e-77, respectively, Figure 3D). After thermal stresses, the strength of H3K27ac tends to increase to a significantly larger extent at top up regulated genes than that in top down regulated genes (p=9.08e-64 and 5.90e-27 at CS and HS, respectively, Figure S5D). Together, the chromosome compartment, accessibility, and epigenetic landscape are largely stable in the response to thermal stress, thus, the transcriptional changes do not directly link to the changes of the three. However, the strength of three in NM condition are clearly distinguished between the transcriptionally up and down regulated gene, suggesting that the epigenetic preconditions may critical to how cell deal with the thermal stresses.

### TAD structure are stable after the thermal stresses

We next asked if there is also genome widely changes of the TADs in response to the thermal stresses, as observed in *Drosophila* (Djekidel et al., 2015). By applying a recently developed algorithm deDoc to our Hi-C data (Li et al., 2018), we identified 2107, 2207 and 1967 TADs in NM, HS and CS, at 10Kb resolution respectively. The accuracy of TADs boundaries was evidenced by the enrichment of CTCF, and SMC3 ChIP-seq peaks. There are 47.84%, 48.59% and 43.11% TAD boundary regions, i.e., the 20kb flanking region of the boundaries, having at least one CTCF ChIP-seq peaks in CS, NM and HS, respectively. Similarly, there are 40.29%,28.05% and 44.65% TAD boundary regions having at least one SMC3 peaks, respectively. Moreover, most of, e.g., about 98.17%, 99.49% and 92.33%, TAD boundaries with at least one SMC3 also has at least one CTCF binding peak in CS, NM and HS, respectively (Figure 4A and B) and SMC3 (Supplemental Figure S7).

**Figure 4.**
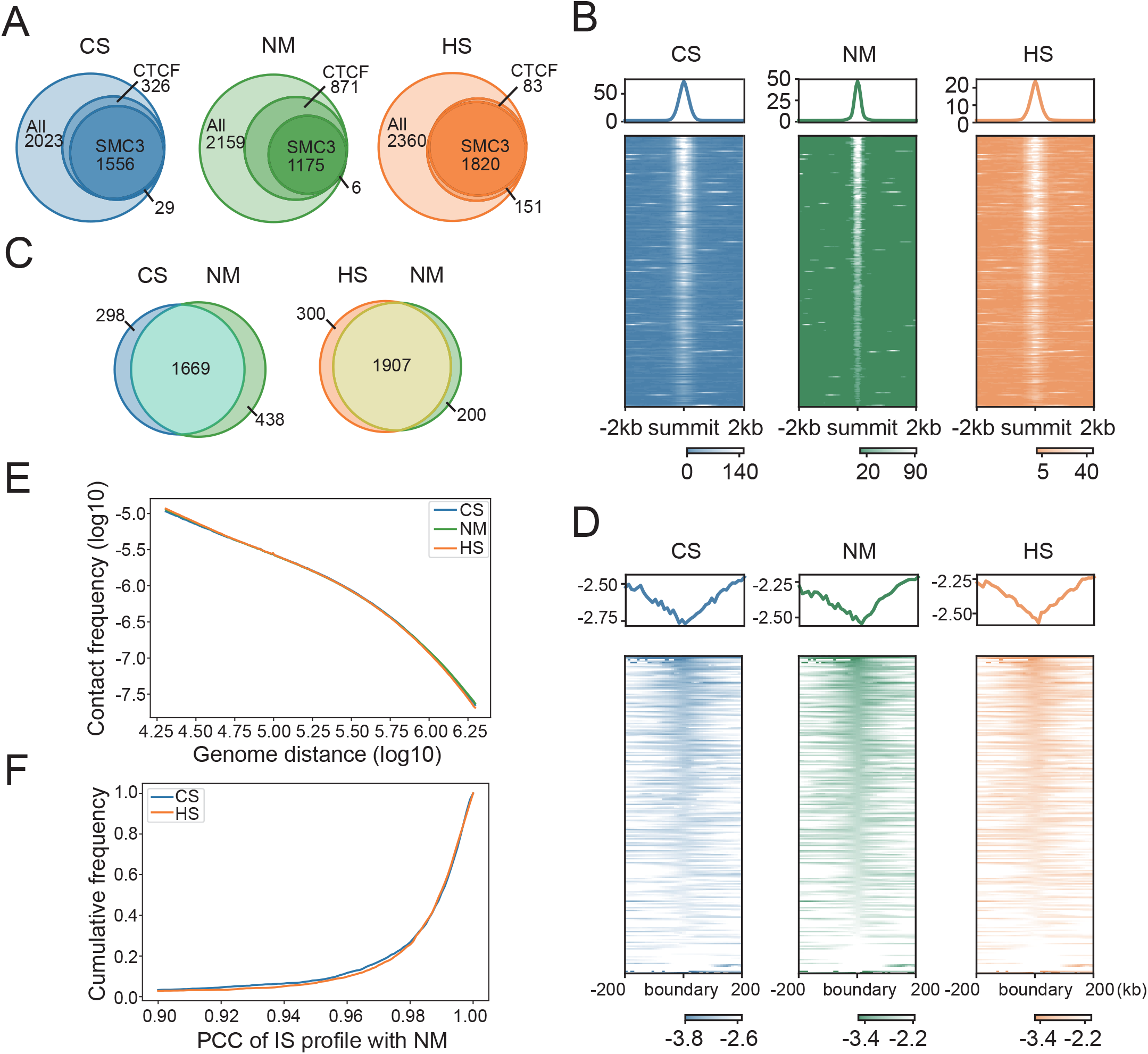
The TAD structures are stable in response to the thermal stresses. (A) The venn diagram shows the overlapping of the TAD boundaries and ChIP-seq peaks of CTCF and SMC3 in the three thermal conditions. (B) Heatmap of CTCF binding affinity at the TAD boundaries in each conditions. The peaks were aligned by their summits, and the cumulative distribution curve were shown on top of each heatmap. (C) The venn diagram shows the overlapping of the TADs between NM and the thermal conditions. (D) Heatmap of insulation score at the TAD boundaries in each conditions with binsize = 10kb around the TAD boundaries at NM. Tracks are centered at the boundary loci and the cumulative distribution curve were shown on top of each heatmap. (E) Contact frequency decay curves drawn with intra-TAD reads only. (F) The cumulative frequencies of the PCC of insulation score profiles in the NM TADs regions between the conditions of NM and the thermal stresses.

We found that the TAD structure are largely stable upon the thermal stresses. First, the alterations of TAD arrangement are minor after the thermal stress. Although the TAD lengths (median 1.06Mb in NM) are slightly changed after CS (median = 1.15Mb) or HS (median = 1.03Mb, Mann Whitney U test p=2.19e-5 and 0.0040, respectively, Figure S8), the TAD can be aligned to each other, i.e. about 84.85% and 86.41% TADs detected at CS and HS can be aligned to at least one TAD in NM with more than 50% overlapping in length (Figure 4C). This alignment rate is comparable to 87%, the alignment rate between two Hi-C replicates in NM. Second, the insulation score (IS) profiles around the TAD boundaries in NM are indistinguishable from the profiles of other two conditions (Figure 4D). Third, the internal structure of TADs does not change after thermal stress. The regressed p(s) curves using intra-TAD Hi-C read only are almost identical between the three conditions (JSDs < 10-4 for both CS and HS, Figure 4E). Correspondingly, the intra-TAD Hi-C contact maps are almost identical between conditions, i.e., comparing with NM, the mean and confidential interval of PCCs of intra-TAD Hi-C contact matrices are 0.9791 (0.9714 to 0.9886 for two sided 95% confidence interval) and 0.9893 (0.9825 to 0.9969) in CS and HS, respectively (Figure S8B). Furthermore, the insulation score (IS) profile within the TAD regions are highly correlated between NM and both CS and HS, i.e, the PCC are 0.9693 (0.8763 to 0.9992 for two sided 95% confidence interval), and 0.9704 (0.8839 to 0.9997), for CS and HS, respectively, Figure 4F). Together, the 3D genome architecture of human cells is largely stable at TAD level under the thermal stress conditions.

### The transcription regulation and alteration of TADs are uncoupled in response to the thermal stresses

Although, the TAD structure were almost stable under thermal stress conditions, there are a small fraction of altered TAD, e.g. fused, split or resized. We wonder whether those altered TADs were associated with the transcriptional response, particularly for those up-regulated genes, as enhancer hijack is recognized as a mechanism to gene regulation in mammals (Symmons et al., 2016). Because the genes in a same TAD prone to be regulated in same direction, i.e., being up- or down-regulated simultaneously (Supplemental Text), we define and identified TADs’ reorganization and direction of transcriptional response in both CS and HS (See Methods for the definition, **Table S2**). There is little correlations can be seen between TAD’s alteration and its transcript regulation changes in either CS or HS (p=0.1486 and 0.8198 in CS and HS, respectively, χ^2^ test, Table S2).

We further asked if gene expression changes were associated with the alteration of TADs in response to the thermal stresses. First there are minor enrichment for expression changed genes in those altered TADs, i.e., p=0.0640 and 0.0094, χ^2^ test, in CS and HS, respectively (Table S3). By assuming the intra-TAD enhancers are the dominative regulators of gene expression, we assessed the influence of TAD alteration to gene expression changes. Alteration of a TAD boundary may let some external enhancers being included into this TAD (i.e., enhancer gain), or some internal enhancers being excluded out this TAD (i.e., enhancer loss). We found that gain or loss enhancers dose not significantly affect the chance of being differentially expressed for the genes in response to both CS and HS, unless gain or loss more than 90% of its intra-TAD enhancers (Figure 5A). There are 283 (13.43%) and 259 (12.29%) out of 2107 TADs in NM have 90% enhancers changed in CS and HS, respectively. This results suggested that the enrichments for expression changed genes in the TADs that experienced dramatic alterations. Second, we further considered the activity of cis-regulatory elements (CRE), i.e., promoters and enhancers, to assess the influence of TAD alternation to gene expression changes. We reason that, if the alteration of TAD structure substantially affects gene expression, one would expect that the activity of intra-TAD CREs in one condition should be a better predictor to the gene expression level in the same condition than the intra-TAD CREs does in other conditions. To test this prediction, we developed a simple linear regression model using activity of promoters and enhancers as predictors. In both CS and HS, the model well catches the gene expressions with the TADs annotated in the corresponding conditions (R^2^=0.413, 0.670 and 0.618 in NM, CS and HS, respectively, Figure 5B), and both promoter and enhancer contributes significantly to the prediction, i.e., having nonzero coefficients (Table S4). However, when we applied the model scheme to predict gene expression level at the thermal stressed conditions using the TAD annotation at NM, we could not reject the hypothesis that the predictions are equal (p=0.2918 and 0.0842 in CS and HS, respectively, two sided binomial test, Figure 5B, Table S4). Last, we checked if the model works more condition specific for those dramatically altered TADs. Indeed, if the similarity of TADs becomes too low, the native TADs annotation is essential to have accurate predication (Figure 5C). Together, our result suggested that the minor and moderate alterations TADs have little influence to gene expression changes, dramatic TAD reformation may affect gene regulation in response to the thermal stresses.

**Figure 5.**
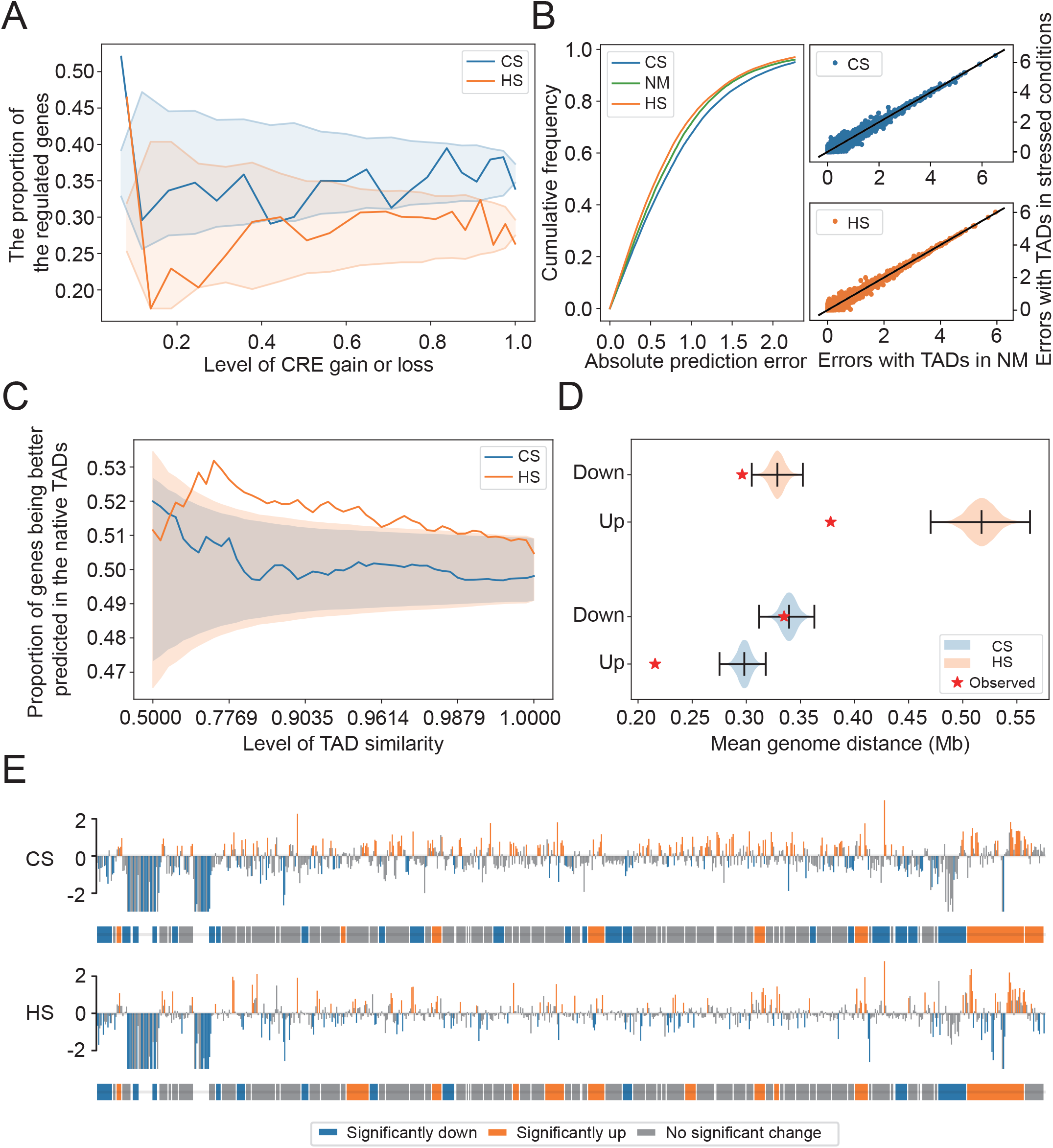
The transcriptional change and alteration of TADs are uncoupled in response to the thermal stresses. (A) Gain or loss CRE is largely uncoupled to gene transcriptional changes. The solid curves represent the cumulative average proportion of regulated genes over the level of gain or loss of CRE, in response to thermal stresses. The shadow represents 0.95 confidence interval of binomial test, with the total regulated genes as control. (B) Left panel: the cumulative distribution of prediction errors of the linear models of gene expressions; Right panels: comparisons of prediction errors of the models with TAD annotations in NM and thermal conditions. (C) The solid curves represent the cumulative distribution of proportion of the genes that being better predicted in the TADs of thermal stressed conditions than in NM, over the similarity of TADs between the conditions. The shadow represents 0.95 confidence interval of binomial test, controlled with the proportion 0.5. (D) The genome distances of the neighboring regulated genes with the same regulation direction. The stars represent the observation in our data, and the violin plots represents the distribution of control with randomly shuffled regulation direction labels. (E) The distribution of the regulated genes and TADs throughout the whole chromosome 8. The vertical bars represent the log2 fold changes of gene expression, the horizontal bars represent the regulated TADs. All the intergenic regions were excluded from this plot.

### Transcription regulation in response to the thermal stresses are beyond TAD level

TADs were thought be the functional units in hormone treatment (Le Dily et al., 2014) and hyperosmotic shock (Amat et al.). We examined if genes in a same TAD tend to have coordinated response to thermal stress. Indeed, there are more TADs that can be assigned a uniformed transcription regulatory direction than random expectation (see Method and Table S2, p<e-4 for all conditions, permutation test). This results verified the transcriptional unities of TADs in response to the thermal stresses, which, however, raises an immediate question, why alteration of TAD structure has little effect to gene expression changes? We speculated that there may exists transcriptional regulatory domains, i.e., the genome regions consist of multiple TADs that have uniformed regulatory directions. If this speculation were correct, we would expect correlated regulation direction of neighboring TADs. Indeed, there are observable significant positive correlated regulation direction of neighboring TADs (p = 0.0143 and 0.0295 in CS and HS, respectively, χ^2^ test, Figure 5D). We further examined if the regulated genes are non-randomly distributed in the whole genome. The mean genome distance of up regulated genes to its nearest up regulated neighbor is significantly smaller than randomly expected (p<1e-4 in both CS and HS, permutation test), it is also true in HS (p<1e-4, Figure 5D and E). Together, our results suggested that the TADs and genes with similar transcription response to thermal stress prone to cluster in the genome.

### The thermal stresses responded TADs are spatially clustered

It is an immediate question to ask if the regulated gene and TADs were spatially clustered, since they were already clustered in genome (Figure 5D and E). To show this clustering in physical space, we modeled the chromosomes structure using TADs as monomers (Lesne et al., 2014). The contact matrices of modeled chromosomes and Hi-C are highly correlated, i.e., the mean PCC are 0.69, 0.72, and 0.67 for NM, CS and HS, respectively (Figure S9). As expected, the modeled chromosomes structure in NM are rather similar to the model in thermal stress conditions (Figure 6A and Figure S9B and C), i.e., the mean SpCC of TAD-TAD distances are 0.96 and 0.97 between NM and CS and NM (Figure S9B and C), respectively. All those similarity implies that the models have captured major characters of chromosomes physical structures. Then, we compared the average distances of nearest neighboring up regulated TADs, and found they are significantly smaller than randomly expected (p = 0.0207 and 0.0924, as the regulation after CS and HS, respectively, permutation test). The same pattern also been observed for down regulated TADs (p = 0.0351 and 0.0875, as the regulation after CS and HS, respectively, permutation test). Further, the regulation directions of spatially nearest neighboring TADs are also significantly positively correlated (Figure 6B, p = 0.0170 and 6.74e-5 as the regulation after HS and CS, respectively, χ^2^ test). Together, the physical model showed a clear spatial clustering of the TADs with the same regulation direction in NM, further suggested the prewired plan for the thermal stress response in the genome 3D architecture.

**Figure 6.**
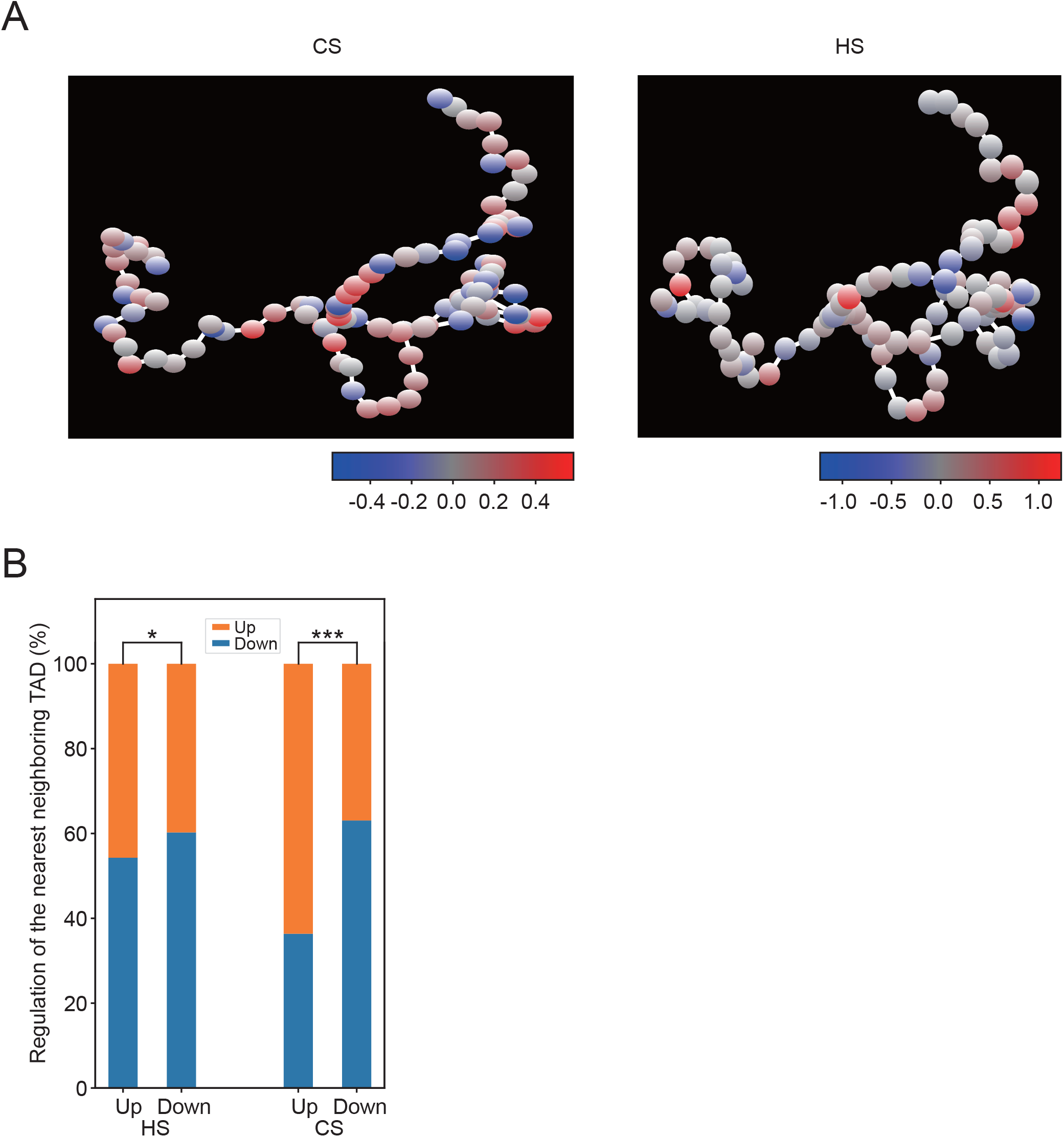
Regulated TADs are spatially clustered. (A) An example of simulated physical structure of chromosome 6 in the thermal stressed conditions. Each monomer represents a TAD, and color of red and blue represent the regulation direction of up and down as defined by log2 fold change of Pol II ChIP-seq signal, respectively. The brightness of colors quantifies the fold changes. (B) The regulation directions of spatially nearest neighboring TADs are positively correlated. The stacking bars represent the percentage of regulation directions of the nearest neighbor TADs for each TAD groups. *: P< 0.1, *** P < 0.001, χ^2^ test.

### The thermal stresses responded genes are associated with high binding affinity of CTCF/cohesin at NM while the chromatin loops are largely stable

Because the distal *cis-*elements (DCE), e.g., enhancers, regulates gene expression via DCE-promoter interactions (Palstra et al., 2003), we asked whether the transcription changes in response to the thermal stresses is regulated by those chromatin loops. If this is true, we would expect the changes of binding affinities of CTCF or cohesin at the regulated promoters or DCEs, as the two are believed key proteins in loop formation (Merkenschlager and Nora, 2016). To test this prediction, we performed ChIP-seq on CTCF, SMC3 and H3K27ac in 3 conditions. Indeed, the binding affinity of the two architectural proteins are significantly changed at the promoters or their DCE after thermal stress (Figure 7A). For promoters, there are 1094 out of 18480 and 1251 promoters (TSS-2Kb to TSS+2Kb) were found have CTCF binding affinity changed (FDR<0.05 (Stark and Brown, 2011)) in CS and HS, respectively, and 1863 and 3159 promoters were found have SMC3 binding affinity changed in CS and HS, respectively. For DCEs, we defined the DCE loci as merged distal ChIP-seq peaks of H3K27ac in the three conditions, i.e., the peaks do not overlap with any promoter region (TSS-2Kb to TSS+2Kb), and the peak height as the activity of the DCE. Similarly, there are a large number of DCEs responded to thermal stress with changed activity, and CTCF and SMCs binding affinity were also found changed in response to thermal stress (Table S6). We further examined whether the changes of CTCF/cohesin is coincident with the changed activity of the promoters or DCEs. The changes of gene expression and the changes of CTCF binding affinity were found significantly correlated, i.e, 40.56% (428 out of 1055) and 32.12% (388 out of 1208) of genes with changed CTCF binding at promoters are also differentially expressed in CS and HS, respectively (p = 5.09e-5 and 3.37e-4, in CS and HS, respectively, χ^2^ test). In addition, for those regulated genes with changed CTCF binding affinity, the CTCF binding affinity change is positively correlated with gene regulation changes (Figure 7B, p = 7.37e-25 and 7.64e-3 in CS and HS, respectively, χ^2^ test). This correlation was also found with SMC3, i.e., there are 37.47% (670 in 1788, p = 0.012) and 31.63% (966 in 3054, p = 5.80e-8) promoters with changed SMC3 binding affinity also experienced gene expression changes in CS and HS, respectively, and the SMC3 binding affinity change directions are also positively correlated with the gene regulation directions (p = 1.53e-41 and 1.02e-12 in CS and HS, respectively, Figure 7B). For DCEs, there are 1846 out of 25563 and 9258 DCEs changed activity in HS and CS, respectively. In those DCEs with changed activity, the correlation was also found between the CTCF/cohesin binding and the activity of DCEs in HS, i.e., CTCF (p = 1.45e-14 and 0.0115, χ^2^ test, Figure 7C) and SMC3 (p=1.13e-13 and 3.04e-5, Figure 7C), in CS and HS, respectively. This correlation encouraged us to further test if the changes of gene express may associate with the alternation of chromatin loops, particularly in the loops related to the promoters or DCEs.

**Figure 7.**
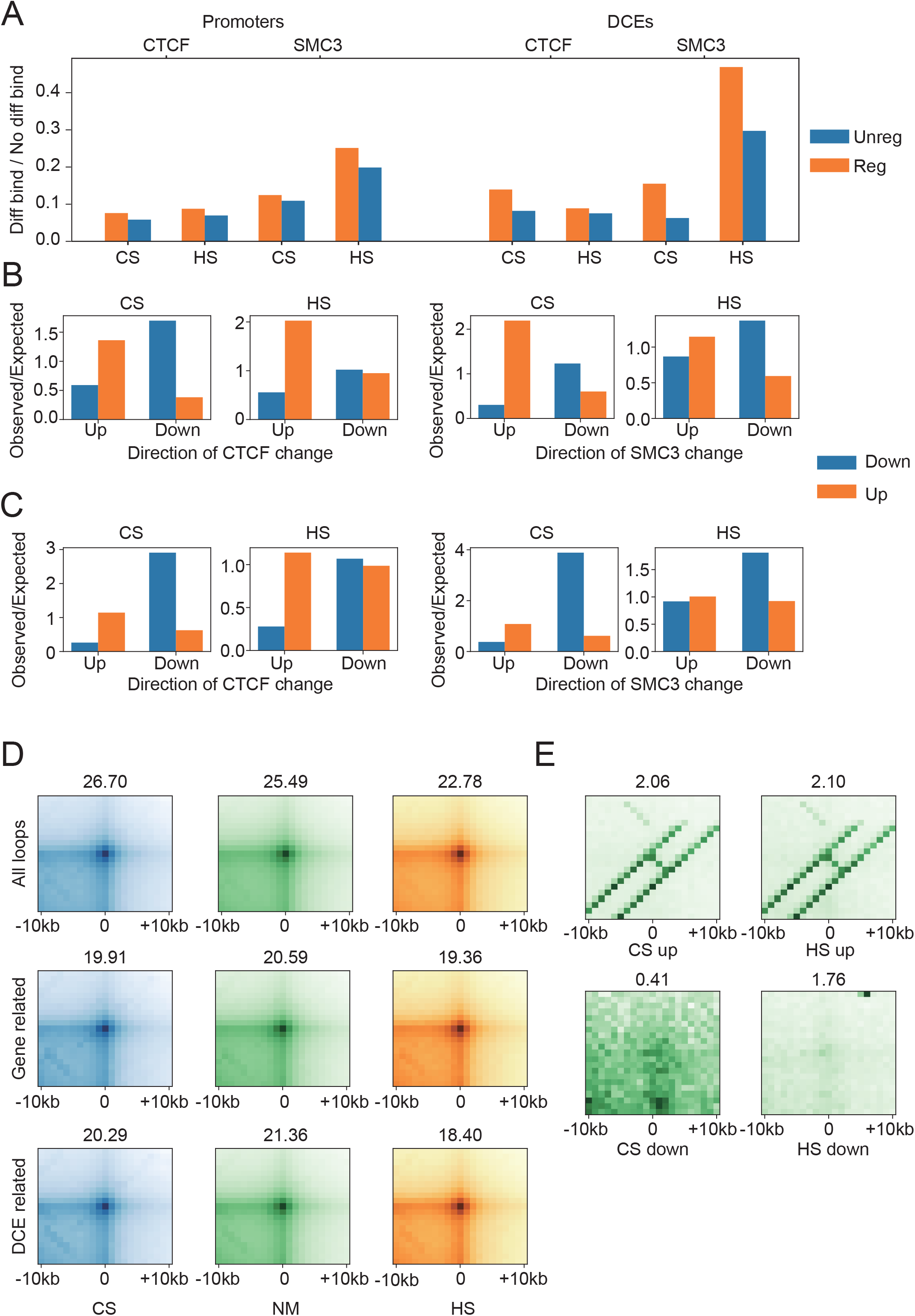
The changes of transcription and binding affinity of CTCF/SMC3 are associated in response to the thermal stresses with stable chromatin loops. (A) The binding affinities of CTCF and SMC3 are prone to changes in the activity changed regulatory elements. The bars above and below the x-axis represent activity changed and unchanged regulatory elements, respectively. The bar heights represent the ratio number of binding affinity changed to unchanged TFs in each group. The change of TF binding affinity are correlated with the change of activities of promoter (B) and DCEs (C). Each bar represents the Observed / expect number of regulated elements grouped by the changes of element activity. The change of binding affinity was color coded. The expect values are calculated using a two-way contingency table. The heatmaps show the APA analysis in (D) and (E), the APA scores were marked above each heatmap. (D) APA results of the HiChIP identified loops that the loop anchors overlapping to CTCF peaks, promoters and DCEs at the three conditions are showed in upper, middle and bottom rows, respectively. (E) The APA analysis at NM of hypothetic randomly paring loops within activity increased and decreased super enhancers are showed in upper and bottom rows, respectively.

If the changes of gene express were regulated by the alternation of chromatin loops, one shall expect the correlated changes between gene expression and loop strength. To test this speculation, we generated CTCF HiChIP data in NM, and called chromatin loop with hichipper package (Lareau and Aryee, 2017). In total, there are 21884 CTCF related loops were identified, in which 13457 having at least one anchor within the 20Kb region (TSS-10Kb, TSS+10Kb) of any TSS, we term them as promoter related loops, and 12570 having at least one anchor within the 20Kb region (middle of DCE-10Kb, middle of DCE+10Kb) of any DCE, we term them as DCE related loops. However, we failed to find significant changes of, either in promoter or DCEs related, chromatin loops using aggregated peak analysis (APA) on Hi-C data. For all the chromatin loops identified in NM, the accumulative strength is almost identical between NM, HS and CS, as, the APA scores are 25.49, 22.78, and 26.70, respectively (Figure 7D). Further, it is also true for the loops related to promoters or DCEs (Figure 7D). Last, even for loops related to up- and down-regulated promoters (Figure S10A) or loops related to the activity changed DCEs (Figure S10B), do not substantially change the strength in CS or HS. These observations imply that the chromatin loops are not only overall largely stable, but also remains stable in regulated promoters and DCEs, in response to thermal stresses.

Given the observed correlations between binding affinity of CTCF/SMC3 and regulated promoters/DCEs, while the chromatin loops remain largely stable, we speculated that the chromatin loop in NM may prewired the plan for thermal stress._ To test this speculation, we compare the chromatin looping strength and the regulation direction of the genes in response to thermal stress. Indeed, we found that the strength of promoter related looping in NM condition is strongly associated with how those genes response to thermal stress. For any given set of loops, the accumulative strength of looping was assessed by the aggregated peak analysis (APA) with the Hi-C data in the condition studied. We compared the looping strengths of up- and down-regulated genes in NM. The accumulative strength of loops related to up-regulated genes are always stronger than that of down regulated genes, i.e, the APA scores are 15.35 and 10.56 for 4176 and 3298 up and down-regulated gene related loops, respectively, in response to CS (Fig. S10A). It is also true for the regulated genes in response to HS (Fig. S10A). For DCEs, the enhancement of stronger chromatin loops flanking the activity increased DCE were also been observed in compare with the activity decreased DCEs, by the APA analysis, i.e., the APA enrichment score of CS up and down of DCE(+/-10k) related loops are 17.67 and 9.41, respectively, in NM (Fig. S10B). Similar results are also observed in HS regulated DCEs (Fig. S10B). Last, this enrichment can also be seen when the activity changes defined by the ChIP-seq data of Pol II, i.e., 7490 and 9564 out of 25563 DCEs with significant activity changes are observed after CS and HS, respectively. The Pol II determined regulation direction of DCEs also significantly positively correlated with both direction of changes of CTCF (p=1.82e-6 and 6.40e-6 in CS and HS, χ^2^ test, Figure S11A) and SMC3 (p = 3.09e-10 and 1.86e-16, Figure S11A). The APA enrichment score of up regulated DCEs related loops (14.88 and 18.31 in CS and HS) is also much larger than that of down regulated DCEs related loops (10.54 and 10.05, Figure S11B).

Activated super enhancers (SEs) after thermal stresses also show stronger spatial contiguity in NM compared as repressed ones. Among the 742 annotated SEs in dbSUPER database (Khan and Zhang, 2016), 99 and 90 in CS and 162 and 152 in HS are defined as down and up regulated at FDR = 0.05, respectively, using the H3K27ac data. The proximity of SEs in each category was measured by APA score of random paired SEs in NM. For both CS and HS, the APA enrichment score between up regulated SEs (2.06 and 2.10 in CS and HS, respectively) are significantly larger than those between down regulated SEs (0.41 and 1.76 in CS and HS, respectively, Figure 7E). Taken together, the strong correlated between the transcription response to the thermal stresses and the strength of chromatin loops in NM condition suggested prewired chromatin loops pattern may largely determine how cells response to the thermal stress, while the chromatin loops may remain stable per se after the stress.

## Discussion

In this study, we explore how cells responses to thermal stresses with the changes of transcription activity, epigenetic marks and chromatin organization. Thousands of genes were found changed the expression level, and surprisingly, conventional chromatin organization characteristics of nuclei – compartments, TADs and interactive loops – are largely stable in response to the thermal stresses, even in the loci containing the most regulated genes. However, in *Drosophila*, short term HS can substantially remodel chromatin structure (Li et al., 2015). These results suggest that warm-blooded animals may have specific mechanisms to allow them respond rapidly to environmental temperature changes without dramatic changes in chromatin structure to survive thermal stress. During the preparation of our manuscript, Judhajeet et al. released an article in the bioRxiv with an observation of stable chromatin 3D organization during HS also (Ray et al., 2019), but how cell response to CS remains unreported. Last, we showed evidences that the plans for the stressful conditions may already encoded in the spatial architecture of genome in normal condition.

Although chromatin 3D architecture is robust, it also needs to be flexible enough to allow dynamics. The hierarchical 3D chromatin structure undergoes remodeling during several processes, such as cell cycle, differentiation, and early embryonic development (Hu et al., 2018; Hug and Vaquerizas, 2018; Nagano et al., 2017). In response to stimulus, it has been shown that after hormone treatment the borders of the ~2000 TADs in T47D breast cancer cells are largely maintained. Intra-TAD interactions exhibited gene activity-related changes with repressed TADs more likely to lose intra-TAD interactions and become more compacted (Le Dily et al., 2014). The discrepancy of cell response towards hormone treatment and thermal stresses is therefore poses an interesting question on possible alternative mechanisms to chemical and physical stimulus(Chen et al., 2017; Kim et al., 2018), and need to be further investigated.

The stability of the chromatin 3D structure and the similarity of cellular response to CS and HS suggest that there are some pre-existing adaptive mechanisms for cells. This remind us the pre-existing chromatin looping (Jin et al., 2013) A previous study showed that TNF-α responsive enhancers are unexpectedly found already in contact with their target promoters prior to stimulation and such pre-existing chromatin looping also exists in other cell types with different extra-cellular signaling (Jin et al., 2013). No changes are detected in chromatin loops upon HS or CS in our study even in the most regulated genes and this is in agreement with the conclusion of this article. This imply that the pre-existing looping may be a widely used mechanism for cells in respond to environmental stimulus. Identifying and understanding the code of the prewired design of chromatin structure for thermal stress, and possibly for other environmental stresses, is critical in synthesis of environmental tolerant systems in the future.

Previous studies showed that the release of promoter-proximal paused RNA polymerase II into elongation is a critical step determining gene’s response to HS stress (Vihervaara et al., 2018). We showed evidences to support that the same inhibition of pause-release of pol II from down-regulated gene’s promoters exists upon CS, implying this may be a common mechanism for cells to rapidly respond to thermal stimulus. In addition, our results demonstrated that the transcriptional reprogramming does not associate with the global chromatin conformation changes in the responses to thermal stress. This is in line with previous studies demonstrated that transcriptional profile was only modestly affected during marked reorganization of chromosomal folding in conditional depletion of cohesin or CTCF (Nora et al., 2017; Rao et al., 2017; Schwarzer et al., 2017). Thus, at least in the condition of the thermal stresses, and maybe in more stress response conditions, changes in chromatin 3D structure and in transcription can be less coupled.

## Method

### Cell Culture and heat shock treatment

K562 erythroleukemia cells were maintained at 37 °C in a humidified 5% CO_2_ atmosphere and cultured in RPMI medium (Sigma) containing 10% (vol/vol) FCS, 2mM L-glutamate, and streptomycin/penicillin. The K562 cells were obtained from ATCC and were tested to be mycoplasma free. To avoid influence of freshly added media, the cells were expanded 24h prior to thermal treatment. For heat shock or cold shock, the growing cells were placed in water bath at 42 °C or 4°C for 30 min.

### In situ Hi-C

In situ Hi-C was conducted according to (Ke et al., 2017). Briefly, samples were fixed with a final concentration of 1% formaldehyde and quenched with 0.125M glycine. Cells were lysed in ice-cold Hi-C lysis buffer (10 mM Tris-HCl pH 8.0, 10 mM NaCl, 0.2% Igepal CA630, 1x protease inhibitor cocktail) for 15 min. Pelleted nuclei were washed once with 1x NEBuffer 2 and incubated in 0.5% sodium dodecyl sulfate (SDS) at 62°C for 5 min. After incubating, water and Triton X −100 were added to quench the SDS. MboI restriction enzyme (NEB, R0147) were added and chromatin was digested. Biotin-14-dATP was used to mark the DNA ends followed by proximity ligation in intact nuclei. After crosslink reversal, samples were sheared to a length of ∼300 bp, then treated with the End Repair/dA-Tailing Module (NEB, E7442L) and Ligation Module (NEB, E7445L) following the operation manual. Biotin-labeled fragments were pulled down using Dynabeads MyOne Streptavidin T1 beads (Life technologies, 65602). The Hi-C library was amplified for about 10 cycles of PCR with Q5 master mix (NEB, M0492L) following the operation manual. DNA was then purified with size selection, quantified and sequenced using an Illumina sequencing platform.

### ChIP-seq library preparation

ChIP-seq was conducted according to (Zhu et al., 2019) with few modifications. The cells were cross-linked with a final concentration of 1% formaldehyde followed by quenching with glycine. Cells were lysed with lysis buffer (0.2% SDS;10 mM Tris - HCl, pH 8.0; 10 mM EDTA, pH 8.0; proteinase inhibitor cocktail) and sonicated to fragments about 300 - 500 bp (Bioruptor, Diagenode). Dynabeads Protein A was washed twice with ChIP Buffer (10mM Tris-HCl pH7.5, 140mM NaCl, 1mM EDTA, 0.5mM EGTA, 1% Triton X-100, 0.1% SDS, 0.1% Na-deoxycholate, Cocktail proteinase inhibitor) and was incubated with antibody at 4℃ for 2-3hours. The fragmented chromatin was transferred to the bead-antibody complex tubes and rotated at 4 °C overnight. The beads were washed once with low salt buffer (10mM Tris-HCl pH7.5, 250mM NaCl, 1mM EDTA, 0.5mM EGTA, 1% Triton X-100, 0.1% SDS, 0.1% Na-deoxycholate, Cocktail proteinase inhibitor) and twice with high salt buffer (10mM Tris-HCl pH7.5, 500mM NaCl, 1mM EDTA, 0.5mM EGTA, 1% Triton X-100, 0.1% SDS, 0.1% Na-deoxycholate, Cocktail proteinase inhibitor). After crosslink reversal, library was constructed following Illumina’s instructions.

### HiChIP

HiChIP was modified from (Mumbach et al., 2016). 15 million crosslinked cells were lysed using Hi-C lysis buffer. Nuclei pellet was incubated in 0.5% SDS at 62°C for 10 minutes. Triton X-100 was added to quench the SDS. MboI (NEB, R0147) was used to digest chromatin overnight. Biotin-labelled DNA ends were proximity ligated in intact nuclei. The nuclei were then divided into 10 parts with 1.5 million nuclei each. The nuclear pellet was sonicated in 220 μL Nuclear Lysis Buffer (50 mM Tris-HCl pH 7.5, 10 mM EDTA, 1% SDS, 1X Roche protease inhibitors) using Bioruptor in the following parameters: ON = 30s, OFF = 30s, Duty Cycle = 1 and then clarified by centrifugation for 10 minutes at 12000g at 4°C. Immunoprecipitation and washing were performed identical to ChIP-seq. Cross-links were reversed and DNA was purified with the Agencourt AMPure XP beads (Beckman Coulter, A63881). Ten tubes of DNA were then combined. After end repair and adaptor ligation, biotin-labeled fragments were pulled down and sequencing library was constructed consistent with In situ Hi-C.

### ATAC-seq

ATAC-seq was prepared as previously described with few modifications (Corces et al., 2017). Briefly, 50000 fresh cells were resuspended in 50 μl of ATAC-seq resuspension buffer (RSB; 10 mM Tris-HCl pH 7.4, 10 mM NaCl, and 3 mM MgCl2) containing 0.1% NP40, 0.1% Tween-20, and 0.01% digitonin and incubated on ice for 3 min. After lysis, 1 ml of ATAC-seq RSB containing 0.1% Tween-20 (without NP40 or digitonin) was used to wash nuclei. Nuclei were resuspended in 50μl of transposition mix (10μl 5XTTBL (Vazyme TD501), 5μl TTE Mix V50, and 35μl water) and pipetted up and down 20 times to mix. Transposition reactions were incubated at 37 °C for 30 min in a thermomixer. After the tagmentation, purify sample using the Ampure XP beads. The ATAC-seq library was amplified for 11 cycles of PCR with TAE mix (Vazyme TD501) following the manual. DNA was then purified with size selection, quantified and sequenced using an Illumina sequencing platform.

### Immunofluorescence analysis

About 50,000 cells were dropped on a glass slide and fixed with 3% paraformaldehyde for 20 min. Cells were then permeabilized by 0.5% Triton X-100 for 5 min. After washing with PBS for three times, cells were blocked with 3% BSA in PBS for 1 h. Nuclear counterstaining was performed with DAPI for 5 min at RT. Then mount the coverslip onto glass slides and seal the coverslip with nail polish. Images were acquired at an RPI Spinning Disk confocal microscope with a 100x objective using MetaMorph acquisition software and a Hammamatsu ORCA-ER CCD camera.

### Sequencing reads pre-processing and quality control

The quality of all libraries are assessed using FasqQC. Reads with mean quality score less than or equal to 30 are removed. For ChIP-Seq and Hi-C libraries, the 5’ most 15bp of both read 1 and 2 are clipped out for the low complexity. Adapters are removed by cutadapt and extremely short fragment with length less than or equal to 30bp (for ChIP-Seq and Hi-C library) are then removed.

### ChIP-Seq data analysis

All ChIP-Seq reads are mapped using bowtie2 with very-sensitive configuration. Fragments with both ends uniquely mapped with MAPQ larger than 5 are then extracted using samtools. Duplicates are removed by picard tools MarkDuplicates. Peaks are called using MACS2 callpeak command using q value 0.01(Zhang et al., 2008). Narrow peaks are called for all libraries. Fold enrichment over control signal tracks were built using the command bdgcmp inMACS2. Peaks called from all replicates of each condition are merged according to an irreproducible discovery rate (IDR) threshold of 0.01. Reads are then merged and a final fold enrichment over control track are made for each condition.

### Differential gene expression analysis using PolII ChIP-Seq

The RefSeq gene annotation was taken from UCSC genome browser. Genes in ChrY and ChrM together with those length less than 2000bp were removed. Read coverage of regions between TTS-50Kb and TTS-500bp of each extremely long genes (longer than 150Kb) were calculated in treated conditions and regressed against the same values in normal condition to get the size factor. Differential analysis was then conducted using DESeq2 and the determined size factor. The CPM value of each gene is calculated by analog to the transcription per million (TPM) value in RNA-Seq data.

### Hi-C and HiChIP data processing

Hi-C reads were processed using Juicer pipeline (Durand et al., 2016). Contact reads related with ChrY and ChrM or with MAPQ = 0 were filtered out. The calculation of contact frequency decay curve is conducted using 501 edge points flanking 20Kb to 50Mb exponentially. A/B compartment profile is called by analyzing the first eigevector of KR normalized contact maps at 100Kb resolution (Lieberman-Aiden et al., 2009b). The compartment with higher H3K27ac ChIP-Seq signals were determined as compartment A. TADs were called using deDoc at 10Kb resolutions (Li et al., 2018). For HiChIP data, peaks of CTCF were directly taken from the CTCF ChIP-Seq dataset and loops were called using HiChIPper (Lareau and Aryee, 2017). Aggregated peak analysis is conducted with respect to loops with anchors located farther than 30 bins in 10Kb contact maps.

Distal local ratio (DLR) was calculated using Hi-C data for each loci following the method of (Heinz et al., 2018). DLR is the log2 fold change of number of Hi-C contact reads linking form the loci to more than 3Mb apart, against the number of contacts linking from the loci to between 20Kb and 3Mb apart.

### Definition of TAD remodeling and direction of TAD regulation

We classified TAD remodeling into five types according to (Ke et al., 2017). ‘Merge’ was defined as multiple TADs in NM fused into one TAD in thermal stress. ‘Split’ was one TAD in NM divided into multiple TADs in thermal stress. ‘Vanish’ was one TAD in NM does not overlap any TADs in thermal stress. ‘Stable’ was one TAD in NM overlapped more than 75% with a TAD in thermal stress. TADs not belong to any of the above groups was classified as ‘other’.

The TAD was defined as upregulated if it contains four or more genes and had significantly higher proportion of genes were upregulated compared to the whole genome, while if a TAD contains four or more genes and had significantly higher proportion of genes downregulated compared to the whole genome, it was defined as downregulated. The direction of gene regulation was determined using the fold change of CPM.

**The similarity of two TADs was defined as**

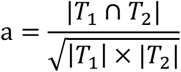

where |T_i_|denotes length of TAD *i*, and |*T*_1_ ∩ *T*_2_| denotes the length of overlapping part of *T*_1_ and *T*_2_.

### Prediction of gene expression according to CRE

The level of gain or loss of CRE was defined as

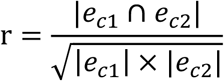

where *e*_*c*_ represents the CREs within the TAD at condition *c*. A smaller number of *r* indicating larger gain or loss CRE between two conditions.

Gene expression level was linearly modeled by the activity of promoter and all the CRE in the TADs. The nascent gene expression level was determined by the CPM described above. The promoter’s activity was quantified by the fold enrichment over control of Pol II ChIP-Seq signal. The activity of CRE was quantified by averaging the fold enrichment of H3K27ac in all CRE located in the same TAD of the query gene. Histone modifications were determined as differential with FDR < 0.01 after the thermal stresses and the direction of the changes were determined by the sign of log2 fold changes.

### Calculation of spatial distance between TADs

In any given condition, for any TAD pair *i* and *j* in the same chromosome, define the contact frequency between *i* and *j* as,

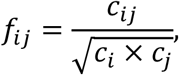

where *c*_*ij*_ denotes the number of reads pairs linking between TAD *i* and *j*, and *c*_*i*_ is the number of intra-chromosome contact read pairs that have and only have one end in TAD i.

The contact frequencies were then transformed into topological distances using the power transformation,

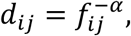

Where *a* denotes power parameter. The adjacent matrix {*d*_*ij*_} defined an undirected graph, and the spatial distances *D*_*ij*_ are calculated as the summation of topological distances in the shortest path between TAD *i* and *j* (Warshall, 1962).

The distance matrix {*D*_*ij*_} are then transformed into the Gram matrix as,

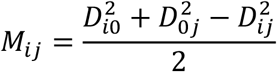

where 0 is an arbitrarily chosen TAD. The three dimensional coordinates for each TAD is calculated by taking the eigenvectors of the largest three eigenvalues of the Gram matrix.

The modeled distances are transformed back into modeled contact frequencies by inversion of the power transform. Power parameter α is estimated by maximize the Pearson correlation coefficient between observed and modeled contract frequencies using Nelder-Mead simplex method initiated at points 1 and 2.

## Reference

Al-Fageeh, M.B., and Smales, C.M. (2006). Control and regulation of the cellular responses to cold shock: the responses in yeast and mammalian systems. Biochem J 397, 247–259.

Amat, R., Böttcher, R., Le Dily, F., Vidal, E., Quilez, J., Cuartero, Y., Beato, M., De Nadal, E., and Posas, F. Rapid reversible changes in compartments and local chromatin organization revealed by hyperosmotic shock. Genome research.

Beer, C., Buhr, P., Hahn, H., Laubner, D., and Wirth, M. (2003). Gene expression analysis of murine cells producing amphotropic mouse leukaemia virus at a cultivation temperature of 32 and 37 degrees C. J Gen Virol 84, 1677–1686.

Carey, H.V., Andrews, M.T., and Martin, S.L. (2003). Mammalian hibernation: cellular and molecular responses to depressed metabolism and low temperature. Physiol Rev 83, 1153–1181.

Chen, D., Toone, W.M., Mata, J., Lyne, R., Burns, G., Kivinen, K., Brazma, A., Jones, N., and Bahler, J. (2003). Global transcriptional responses of fission yeast to environmental stress. Molecular biology of the cell 14, 214–229.

Chen, F.X., Xie, P., Collings, C.K., Cao, K., Aoi, Y., Marshall, S.A., Rendleman, E.J., Ugarenko, M., Ozark, P.A., Zhang, A., et al. (2017). PAF1 regulation of promoter-proximal pause release via enhancer activation. Science 357, 1294–1298.

Chowdhary, S., Kainth, A.S., and Gross, D.S. (2017). Heat Shock Protein Genes Undergo Dynamic Alteration in Their Three-Dimensional Structure and Genome Organization in Response to Thermal Stress. Molecular and cellular biology 37.

Chowdhary, S., Kainth, A.S., Pincus, D., and Gross, D.S. (2019). Heat Shock Factor 1 Drives Intergenic Association of Its Target Gene Loci upon Heat Shock. Cell reports 26, 18–28 e15.

Consortium, E.P. (2012). An integrated encyclopedia of DNA elements in the human genome. Nature 489, 57–74.

Corces, M.R., Trevino, A.E., Hamilton, E.G., Greenside, P.G., Sinnott-Armstrong, N.A., Vesuna, S., Satpathy, A.T., Rubin, A.J., Montine, K.S., Wu, B., et al. (2017). An improved ATAC-seq protocol reduces background and enables interrogation of frozen tissues. Nature methods 14, 959–962.

Creyghton, M.P., Cheng, A.W., Welstead, G.G., Kooistra, T., Carey, B.W., Steine, E.J., Hanna, J., Lodato, M.A., Frampton, G.M., and Sharp, P.A. (2010). Histone H3K27ac separates active from poised enhancers and predicts developmental state. Proceedings of the National Academy of Sciences 107, 21931–21936.

Dixon, J.R., Selvaraj, S., Yue, F., Kim, A., Li, Y., Shen, Y., Hu, M., Liu, J.S., and Ren, B. (2012). Topological domains in mammalian genomes identified by analysis of chromatin interactions. Nature 485, 376–380.

Djekidel, M.N., Liang, Z., Wang, Q., Hu, Z., Li, G., Chen, Y., and Zhang, M.Q. (2015). 3CPET: finding co-factor complexes from ChIA-PET data using a hierarchical Dirichlet process. Genome biology 16, 1–16.

Duarte, F.M., Fuda, N.J., Mahat, D.B., Core, L.J., Guertin, M.J., and Lis, J.T. (2016). Transcription factors GAF and HSF act at distinct regulatory steps to modulate stress-induced gene activation. Genes & development 30, 1731–1746.

Durand, N., Shamim, M., Machol, I., Rao, S.P., Huntley, M., Lander, E., and Aiden, E.L. (2016). Juicer Provides a One-Click System for Analyzing Loop-Resolution Hi-C Experiments. Cell systems 3, 95–98.

Fortin, J.P., and Hansen, K.D. (2015). Reconstructing A/B compartments as revealed by Hi-C using long-range correlations in epigenetic data. Genome Biology 16, 180.

Fujimoto, M., and Nakai, A. (2010). The heat shock factor family and adaptation to proteotoxic stress. The FEBS journal 277, 4112–4125.

Fujita, J. (1999). Cold shock response in mammalian cells. J Mol Microbiol Biotechnol 1, 243–255.

Heinz, S., Texari, L., Hayes, M.G.B., Urbanowski, M., Chang, M.W., Givarkes, N., Rialdi, A., White, K.M., Albrecht, R.A., Pache, L., et al. (2018). Transcription Elongation Can Affect Genome 3D Structure. Cell 174, 1522–1536 e1522.

Hu, G., Cui, K., Fang, D., Hirose, S., Wang, X., Wangsa, D., Jin, W., Ried, T., Liu, P., Zhu, J., et al. (2018). Transformation of Accessible Chromatin and 3D Nucleome Underlies Lineage Commitment of Early T Cells. Immunity 48, 227–242 e228.

Huang Da, W. (2009). Systematic and integrative analysis of large gene lists using DAVID bioinformatics resources. Nature protocols 4, 44.

Hubner, M.R., Eckersley-Maslin, M.A., and Spector, D.L. (2013). Chromatin organization and transcriptional regulation. Current opinion in genetics & development 23, 89–95.

Hug, C.B., and Vaquerizas, J.M. (2018). The Birth of the 3D Genome during Early Embryonic Development. Trends in genetics: TIG 34, 903–914.

Jin, F., Li, Y., Dixon, J.R., Selvaraj, S., Ye, Z., Lee, A.Y., Yen, C.A., Schmitt, A.D., Espinoza, C.A., and Ren, B. (2013). A high-resolution map of the three-dimensional chromatin interactome in human cells. Nature 503, 290–294.

Ke, Y., Xu, Y., Chen, X., Feng, S., Liu, Z., Sun, Y., Yao, X., Li, F., Zhu, W., Gao, L., et al. (2017). 3D Chromatin Structures of Mature Gametes and Structural Reprogramming during Mammalian Embryogenesis. Cell 170, 367–381 e320.

Khan, A., and Zhang, X. (2016). dbSUPER: a database of super-enhancers in mouse and human genome. Nucleic Acids Research 44, D164–D171.

Kim, Y.J., Xie, P., Cao, L., Zhang, M.Q., and Kim, T.H. (2018). Global transcriptional activity dynamics reveal functional enhancer RNAs. Genome research 28, 1799–1811.

Kregel, K.C. (2002). Heat shock proteins: modifying factors in physiological stress responses and acquired thermotolerance. J Appl Physiol (1985) 92, 2177–2186.

Lareau, C., and Aryee, M. (2017). hichipper: A preprocessing pipeline for assessing library quality and DNA loops from HiChIP data. BioRxiv, 192302.

Le Dily, F., Bau, D., Pohl, A., Vicent, G.P., Serra, F., Soronellas, D., Castellano, G., Wright, R.H., Ballare, C., Filion, G., et al. (2014). Distinct structural transitions of chromatin topological domains correlate with coordinated hormone-induced gene regulation. Genes & development 28, 2151–2162.

Lesne, A., Riposo, J., Roger, P., Cournac, A., and Mozziconacci, J. (2014). 3D genome reconstruction from chromosomal contacts. Nat Methods 11, 1141–1143.

Li, A., Yin, X., Xu, B., Wang, D., Han, J., Wei, Y., Deng, Y., Xiong, Y., and Zhang, Z. (2018). Decoding topologically associating domains with ultra-low resolution Hi-C data by graph structural entropy. Nature communications 9, 3265.

Li, L., Lyu, X., Hou, C., Takenaka, N., Nguyen, H.Q., Ong, C.T., Cubenas-Potts, C., Hu, M., Lei, E.P., Bosco, G., et al. (2015). Widespread rearrangement of 3D chromatin organization underlies polycomb-mediated stress-induced silencing. Molecular cell 58, 216–231.

Lieberman-Aiden, E., Van Berkum, N.L., Williams, L., Imakaev, M., Ragoczy, T., Telling, A., Amit, I., Lajoie, B.R., Sabo, P.J., and Dorschner, M.O. (2009a). Comprehensive mapping of long-range interactions reveals folding principles of the human genome. science 326, 289–293.

Lieberman-Aiden, E., van Berkum, N.L., Williams, L., Imakaev, M., Ragoczy, T., Telling, A., Amit, I., Lajoie, B.R., Sabo, P.J., Dorschner, M.O., et al. (2009b). Comprehensive mapping of long-range interactions reveals folding principles of the human genome. Science 326, 289–293.

Lozzio, C.B., and Lozzio, B.B. (1975). Human chronic myelogenous leukemia cell-line with positive Philadelphia chromosome. Blood 45, 321–334.

Lyu, X., Rowley, M.J., and Corces, V.G. (2018). Architectural Proteins and Pluripotency Factors Cooperate to Orchestrate the Transcriptional Response of hESCs to Temperature Stress. Molecular cell 71, 940–955 e947.

Mahat, D.B., Kwak, H., Booth, G.T., Jonkers, I.H., Danko, C.G., Patel, R.K., Waters, C.T., Munson, K., Core, L.J., and Lis, J.T. (2016a). Base-pair-resolution genome-wide mapping of active RNA polymerases using precision nuclear run-on (PRO-seq). Nat Protoc 11, 1455–1476.

Mahat, D.B., Salamanca, H.H., Duarte, F.M., Danko, C.G., and Lis, J.T. (2016b). Mammalian heat shock response and mechanisms underlying its genome-wide transcriptional regulation. Molecular cell 62, 63–78.

Mahat, D.B., Salamanca, H.H., Duarte, F.M., Danko, C.G., and Lis, J.T. (2016c). Mammalian Heat Shock Response and Mechanisms Underlying Its Genome-wide Transcriptional Regulation. Mol Cell 62, 63–78.

Merkenschlager, M., and Nora, E.P. (2016). CTCF and Cohesin in Genome Folding and Transcriptional Gene Regulation. Annual Review of Genomics & Human Genetics 17, 17.

Mumbach, M.R., Rubin, A.J., Flynn, R.A., Dai, C., Khavari, P.A., Greenleaf, W.J., and Chang, H.Y. (2016). HiChIP: efficient and sensitive analysis of protein-directed genome architecture. Nature methods 13, 919–922.

Nagano, T., Lubling, Y., Varnai, C., Dudley, C., Leung, W., Baran, Y., Mendelson Cohen, N., Wingett, S., Fraser, P., and Tanay, A. (2017). Cell-cycle dynamics of chromosomal organization at single-cell resolution. Nature 547, 61–67.

Ni, J.Z., Kalinava, N., Chen, E., Huang, A., Trinh, T., and Gu, S.G. (2016). A transgenerational role of the germline nuclear RNAi pathway in repressing heat stress-induced transcriptional activation in C. elegans. Epigenetics & chromatin 9, 3.

Nora, E.P., Goloborodko, A., Valton, A.L., Gibcus, J.H., Uebersohn, A., Abdennur, N., Dekker, J., Mirny, L.A., and Bruneau, B.G. (2017). Targeted Degradation of CTCF Decouples Local Insulation of Chromosome Domains from Genomic Compartmentalization. Cell 169, 930–944 e922.

Palstra, R.J., Tolhuis, B., Splinter, E., Nijmeijer, R., Grosveld, F., and de Laat, W. (2003). The β-globin nuclear compartment in development and erythroid differentiation. Nature genetics 35, 190–194.

Rao, S.S., Huntley, M.H., Durand, N.C., Stamenova, E.K., Bochkov, I.D., Robinson, J.T., Sanborn, A.L., Machol, I., Omer, A.D., Lander, E.S., et al. (2014). A 3D map of the human genome at kilobase resolution reveals principles of chromatin looping. Cell 159, 1665–1680.

Rao, S.S.P., Huang, S.C., Glenn St Hilaire, B., Engreitz, J.M., Perez, E.M., Kieffer-Kwon, K.R., Sanborn, A.L., Johnstone, S.E., Bascom, G.D., Bochkov, I.D., et al. (2017). Cohesin Loss Eliminates All Loop Domains. Cell 171, 305–320 e324.

Ray, J., Munn, P.R., Vihervaara, A., Ozer, A., Danko, C.G., and Lis, J.T. (2019). Chromatin conformation remains stable upon extensive transcriptional changes driven by heat shock. bioRxiv.

Schwarzer, W., Abdennur, N., Goloborodko, A., Pekowska, A., Fudenberg, G., Loe-Mie, Y., Fonseca, N.A., Huber, W., Haering, C.H., Mirny, L., et al. (2017). Two independent modes of chromatin organization revealed by cohesin removal. Nature 551, 51–56.

Sonna, L.A., Fujita, J., Gaffin, S.L., and Lilly, C.M. (2002). Invited review: Effects of heat and cold stress on mammalian gene expression. J Appl Physiol (1985) 92, 1725–1742.

Stark, R., and Brown, G. (2011). DiffBind: differential binding analysis of ChIP-Seq peak data. R package version 100, 4–3.

Symmons, O., Pan, L., Remeseiro, S., Aktas, T., Klein, F., Huber, W., and Spitz, F. (2016). The Shh topological domain facilitates the action of remote enhancers by reducing the effects of genomic distances. Developmental cell 39, 529–543.

Velichko, A.K., Markova, E.N., Petrova, N.V., Razin, S.V., and Kantidze, O.L. (2013). Mechanisms of heat shock response in mammals. Cellular and molecular life sciences: CMLS 70, 4229–4241.

Vihervaara, A., Duarte, F.M., and Lis, J.T. (2018). Molecular mechanisms driving transcriptional stress responses. Nature reviews Genetics 19, 385–397.

Vihervaara, A., Mahat, D.B., Guertin, M.J., Chu, T., Danko, C.G., Lis, J.T., and Sistonen, L. (2017). Transcriptional response to stress is pre-wired by promoter and enhancer architecture. Nature communications 8, 255.

Wada, Y., Ohta, Y., Xu, M., Tsutsumi, S., Minami, T., Inoue, K., Komura, D., Kitakami, J.i., Oshida, N., and Papantonis, A. (2009). A wave of nascent transcription on activated human genes. Proceedings of the National Academy of Sciences 106, 18357–18361.

Warshall, S. (1962). A Theorem on Boolean Matrices.

Wu, J., Liu, T., Rios, Z., Mei, Q., Lin, X., and Cao, S. (2017). Heat Shock Proteins and Cancer. Trends Pharmacol Sci 38, 226–256.

Yamaguchi, Y., Takagi, T., Wada, T., Yano, K., Furuya, A., Sugimoto, S., Hasegawa, J., and Handa, H. (1999). NELF, a multisubunit complex containing RD, cooperates with DSIF to repress RNA polymerase II elongation. Cell 97, 41–51.

Yang, T., Zhang, F., Yardımcı, G.G., Song, F., Hardison, R.C., Noble, W.S., Yue, F., and Li, Q. (2017). HiCRep: assessing the reproducibility of Hi-C data using a stratum-adjusted correlation coefficient. Genome research 27, 1939–1949.

Zhang, Y., Liu, T., Meyer, C.A., Eeckhoute, J., Johnson, D.S., Bernstein, B.E., Nusbaum, C., Myers, R.M., Brown, M., Li, W., et al. (2008). Model-based analysis of ChIP-Seq (MACS). Genome Biol 9, R137.

Zhu, W., Xu, X., Wang, X., and Liu, J. (2019). Reprogramming histone modification patterns to coordinate gene expression in early zebrafish embryos. BMC genomics 20, 248.

